# PARPi synthetic lethality derives from replication-associated single-stranded DNA gaps

**DOI:** 10.1101/781989

**Authors:** Ke Cong, Arne Nedergaard Kousholt, Min Peng, Nicholas J. Panzarino, Wei Ting Chelsea Lee, Sumeet Nayak, John Krais, Jennifer Calvo, Matt Bere, Eli Rothenberg, Neil Johnson, Jos Jonkers, Sharon B. Cantor

**Affiliations:** University of Massachusetts Medical School; Netherlands Cancer Institute; New York University School of Medicine; Fox Chase Cancer Center

## Abstract

BRCA1 or BRCA2 (BRCA)-deficient tumor cells have defects in DNA double strand break repair by homologous recombination (HR) and fork protection (FP) that are thought to underlie the sensitivity to poly(ADP-ribose) polymerase inhibitor (PARPi). Given the recent finding that PARPi accelerates DNA replication, it was proposed that high speed DNA replication leads to DNA double strand breaks (DSBs). Here, we tested the alternative hypothesis that PARPi sensitivity in BRCA deficient cells results from combined replication dysfunction that causes a lethal accumulation of replication-associated single-stranded DNA (ssDNA) gaps. In support of a gap toxicity threshold, PARPi-induced ssDNA gaps accumulate more excessively in BRCA deficient cells and are suppressed in *de novo* and genetic models of PARPi resistance while defects in HR or FP often lack this correlation. We also uncouple replication speed from lethality. The clear link between PARPi sensitivity and ssDNA gaps provides a new paradigm for understanding synthetic lethal interactions.

## INTRODUCTION

BRCA1- and BRCA2-deficient cancer cells have a synthetic lethal interaction with PARP inhibition (Bryant et al., 2005; Farmer et al., 2005). This landmark finding has led to the use of several PARP inhibitors (PARPi) in the clinic to treat BRCA deficient and other cancers (Ashworth and Lord, 2018; Lord et al., 2014; Mirza et al., 2018). The synthetic lethality is understood to derive from PARPi inducing DNA double strand breaks (DSBs) that persist in BRCA deficient cells due to defects in homologous recombination (HR) (Bryant et al., 2005; Farmer et al., 2005) and fork protection (FP) (Schlacher et al., 2011; Schlacher et al., 2012). Consistent with the DSB inducing model of therapy response in BRCA deficient cells, restoration of HR or FP are associated with chemoresistance (Bouwman et al., 2010; Bunting et al., 2010; Chaudhuri et al., 2016; Edwards et al., 2008; Sakai et al., 2008). However, this model of therapy response is challenged by recent reports indicating that chemotoxic agents do not initially induce DSBs or even pause DNA replication forks (Huang et al., 2013; Mutreja et al., 2018; Zellweger et al., 2015). Indeed, PARPi accelerates DNA replication(Maya-Mendoza et al., 2018). To square this finding with the framework that HR and/or FP defects drive PARPi therapy response, it was proposed that high speed DNA replication ultimately induces DSBs (Maya-Mendoza et al., 2018; Quinet and Vindigni, 2018). However, we recently proposed a competing model in which ssDNA gaps underlie the BRCA deficiency phenotype, and not DSBs and we propose are fundamental to the mechanism-of-action of genotoxic therapies (Panzarino et al., 2019).

Here, we provide evidence that the genotoxic lesion driving PARPi synthetic lethality in BRCA deficient cancer is wide-spread single-stranded DNA (ssDNA) gaps. Gap induction is associated with PARPi potentially due to its role in the repair of ssDNA breaks, processing Okazaki fragments, or regulating replication-fork reversal — a mechanism by which replication forks reverse direction when confronted with replication obstacles (Berti et al., 2013; Ray Chaudhuri et al., 2012; Sugimura et al., 2008),(Caldecott et al., 1996; Hanzlikova et al., 2018; Leppard et al., 2003; Lopes et al., 2006; Maya-Mendoza et al., 2018; Peng et al., 2018; Sogo et al., 2002). Critically, we demonstrate that PARPi gaps are compounded in BRCA-deficient cells, but suppressed in cell lines and tumors with intrinsic, genetic or de novo mechanisms of PARPi resistance. These findings highlight that a key function of the BRCA-RAD51 proteins is to limit replication gaps (Hashimoto et al., 2010; Kolinjivadi et al., 2017a; Kolinjivadi et al., 2017b; Zellweger et al., 2015, Panzarino et al., 2019) and that loss of this function confers therapy response.

## RESULTS

### PARPi generates ssDNA gaps that are elevated in BRCA deficient cells

To test the hypothesis that PARPi generates ssDNA gaps that are elevated in BRCA deficient cells, we analyzed the response of DNA replication forks to PARPi by employing DNA fiber assays. DNA fiber assays monitor replication by incorporating nucleotide analogs into newly-synthesized DNA strands which can then be fluorescently labeled. As such, any replication perturbation can be observed with single-molecule resolution. To determine if PARPi creates ssDNA replication gaps, we employed an isogenic cell system in which BRCA1 was either proficient or deleted by CRISPR-Cas9 in retinal pigment epithelial 1 (RPE1) cells (Noordermeer et al., 2018). As expected the BRCA1 K/O RPE1 cells were sensitive to both the PARPi, olaparib, and the DNA interstrand crosslinking (ICL) agent, cisplatin (**Figure 1A,B)**. Consistent with recent observations, our DNA fiber analysis revealed that following 2-h PARPi treatment, both control and the BRCA1 K/O RPE1 cells displayed longer dual labeled replication tracts (5-Iodo-2’-deoxyuridine (IdU) and 5-chloro-2’-deoxyuridine (CldU)) as compared to untreated (Maya-Mendoza et al., 2018) (**Figure 1C**). Moreover, we confirmed that BRCA1 deficiency lead to greater fork asymmetry that was reduced with PARPi (Maya-Mendoza et al., 2018) (**Figure S1A**). Strikingly, BRCA1 K/O cells had longer dual labelled tracks as compared to control cells (**Figure 1C)** indicating that PARPi sensitivity could correlate with an immediate response of longer or accelerated replication tracts.

**Figure 1:**
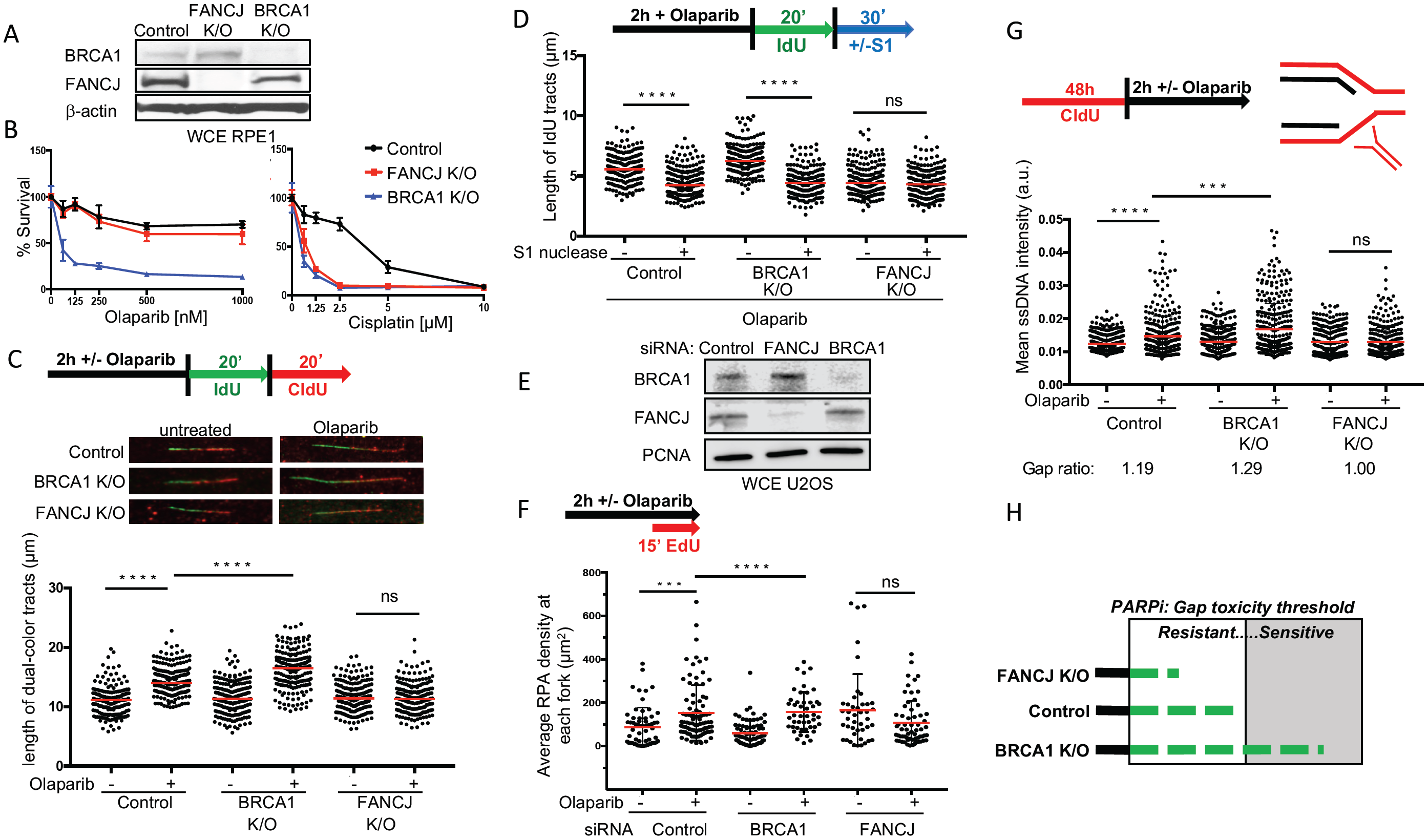
PARPi generates ssDNA gaps that are elevated in BRCA deficient cells but restricted in FANCJ deficient cells. (A) Western blot analysis with the indicated antibodies of lysates from Control, FANCJ K/O and BRCA1 K/O RPE1 cells. WCE, whole cell lysates. (B) Cell survival assays for Control, FANCJ K/O and BRCA1 K/O RPE1 cells under increasing concentrations of olaparib or cisplatin. Data represent the mean percent ± SD of survival for each dot. (C) Schematic, representative images, and quantification of the length of dual-color tracts in indicated cells following olaparib treatment (10μM, 2 hours). (D) Schematic and quantification of the length of IdU tracts with or without S1 nuclease incubation in indicated cells following olaparib treatment (10μm, 2 hours). For (C) and (D), each dot represents one fiber; at least 200 fibers are quantified from two independent experiments. Red bars represent the median. Statistical analysis according to two tailed Mann-Whitney test. (E) Western blot analysis with the indicated antibodies of lysates from U2OS cells expressing siRNA against non-silencing control, FANCJ and BRCA1. (F) Schematic and quantification of average RPA density at each fork in STORM analysis for indicated cells and treatment. Bars represent the mean ± SD. Each dot represents the pair localized RPA density from one nucleus. Statistical analysis according to t test. All p values are described in Statistical methods. (G) Schematic and quantification of mean ssDNA intensity for indicated cells following CldU pre-labeling and olaparib release (10μM, 2 hours). At least 300 cells are quantified from three independent experiments. Bars represent the mean ± SD. Statistical analysis according to t test. The gap ratio is the fold change of ssDNA intensity from treated to untreated. (H) Model in which replication gaps over a toxicity threshold will lead to PARPi sensitivity.

To test the hypothesis that gaps form in the vicinity of the accelerated replication fork, following PARPi treatment, cells were incubated with the S1 nuclease that digests ssDNA regions(Quinet et al., 2017; Quinet et al., 2016). If nascent ssDNA regions are within the labelled replication tracts, S1 nuclease will cut and therefore, shorten the visible CldU replication tract. Following PARPi treatment, labelled nascent DNA tracks appeared shorter for both control and BRCA1 K/O cells when treated with the nuclease (**Figure 1D**). Given that PARPi induces longer initial tract lengths in the BRCA1 deficient cells as compared to control, the similar reduction of tracts lengths following S1 nuclease treatment indicates that PARPi generates a more extensive region of gaps in the BRCA1 K/O cells. To further query the relationship of gaps to individual replication assemblies, we utilized single-molecule localization microscopy (STORM) for direct visualization and quantification of individual replisomes (EdU, MCM6 and PCNA positive sites) and ssDNA bound RPA in the osteosarcoma cell line U2OS (**Figure S1B**). To measure the amounts of incorporated RPA within individual replication domains, we preformed unbiased correlation-based image analyses in control and BRCA1 depleted U2OS cells following PARPi treatment (**Figure 1E**). As compared to the control, the average RPA fork density was greater in the BRCA1 deficient U2OS cells following PARPi treatment (**Figure 1F**). Remarkably, non-denaturing immunofluorescence also revealed that PARPi induced ssDNA gaps genome-wide that were also significantly escalated in BRCA K/O RPE1 cells (**Figure 1G**). Collectively, these findings indicate that PARPi treatment induces gaps that are more pronounced in PARPi sensitive BRCA1 deficient cells.

### Fork acceleration alone does not underlie synthetic lethality caused by PARPi

We previously found that unrestrained replication and gaps caused by loss of the fork remodeler, HLTF were dependent on the BRCA1-associated helicase FANCJ (BACH1/BRIP1), which similar to BRCA1 is a hereditary breast/ovarian cancer and Fanconi anemia gene functioning in HR and FP (Peng et al., 2018),(Cantor et al., 2004; Levitus et al., 2005; Levran et al., 2005; Litman et al., 2005; Nath et al., 2017; Peng et al., 2018; Peng et al., 2006; Sawyer et al., 2015; Suhasini et al., 2013). Similar to BRCA1 K/O cells, we observed that FANCJ K/O cells were sensitive to cisplatin or camptothecin, CPT and maintained similar tracts lengths in unchallenged conditions (**Figure 1B,1C, S1C**). However, in contrast to BRCA1 K/O cells, following PARPi treatment, FANCJ K/O cells did not display PARPi sensitivity, fork acceleration, undergo S1 nuclease cutting, develop RPA rich fork regions or have genomic ssDNA accumulation (**Figure 1C-G, S1D,E**). Given control cells show acceleration and gaps at 10μM PARPi, there could be a gap toxicity threshold that is exceeded in BRCA deficient cells based on a gap ratio (**Figure 1G,H**). However, given that FANCJ is required for PARPi-induced replication acceleration and gaps, either or both could underlie PARPi sensitivity.

To address the role of acceleration and/or gaps in therapy response, we sought to uncouple them. One way to accelerate replication is to deplete the cell cycle regulator, p21 that when combined with PARPi, generates very long replication tracts(Maya-Mendoza et al., 2018) (**Figure 2A-D**). Notably, the lengthening of tracts in p21 depleted U2OS or RPE1 cells did not enhance PARPi sensitivity (**Figure 2E, 2F**). Furthermore, gaps were significantly decreased (**Figure 2G, 2H**). These findings suggest that enhanced fork speed alone is not the cause of PARPi sensitivity (**Figure 2I**).

**Figure 2:**
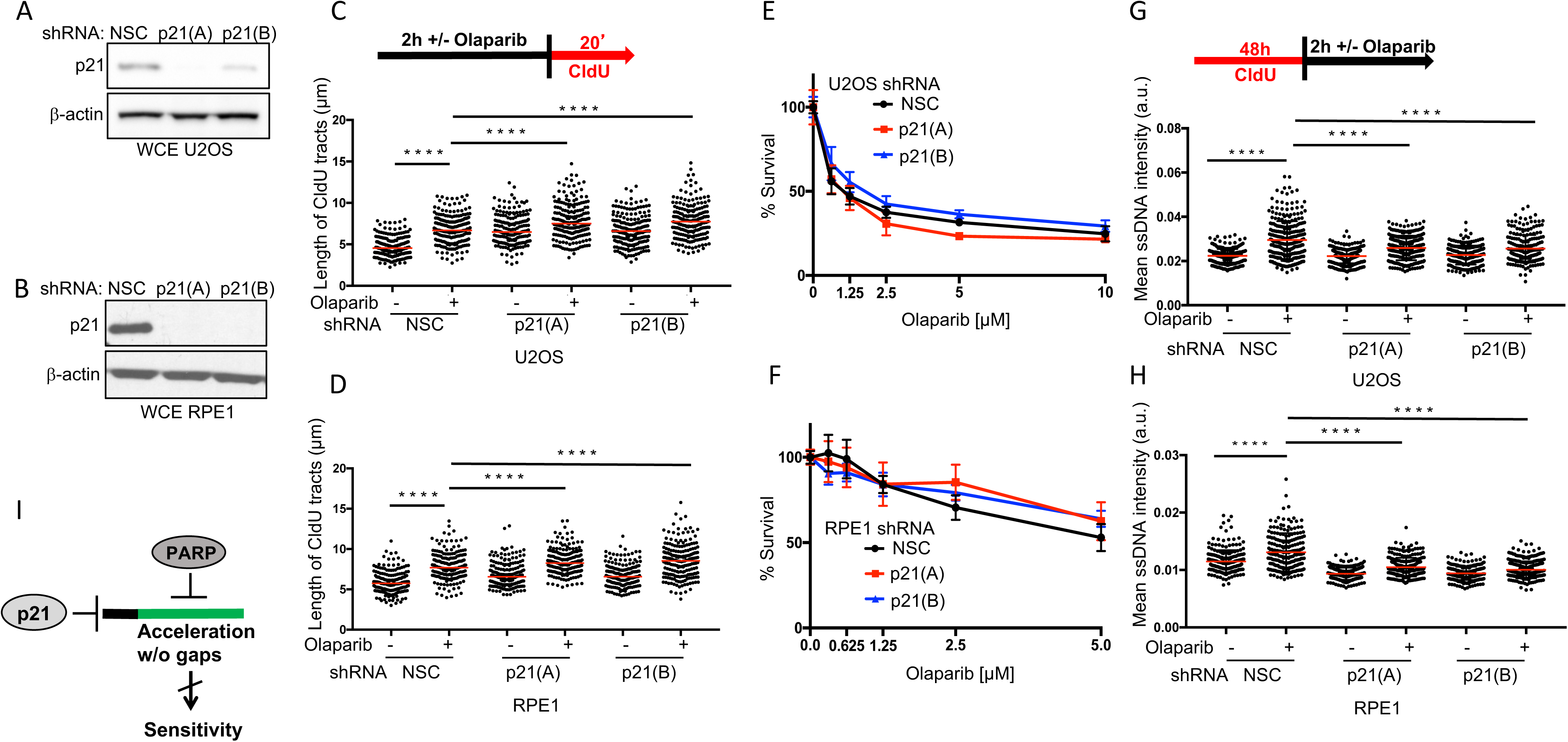
Fork acceleration by p21 depletion does not underlie synthetic lethality caused by PARPi. (A) and (B) Western blot analysis with the indicated antibodies of lysates from U2OS(A) and RPE1(B) cells expressing shRNA against non-silencing control (NSC), p21(A) and p21(B). (C) and (D) Schematic and quantification of the length of CldU tracts in indicated U2OS and RPE1cells following olaparib treatment (10μM, 2 hours). For (C) and (D), each dot represents one fiber; at least 100 fibers are quantified. Red bars represent the median. Statistical analysis according to two tailed Mann-Whitney test. (E) and (F) Cell survival assays for indicated U2OS and RPE1 cells under increasing concentrations of olaparib. Data represent the mean percent ± SD of survival for each dot. (G) and (H) Schematic and quantification of mean ssDNA intensity for indicated cells following CldU pre-labeling and olaparib release (10μm, 2 hours). At least 200 cells are quantified from two independent experiments. Bars represent the mean ± SD. Statistical analysis according to t test. All p values are described in Statistical methods. (I) Model depicting that enhanced fork speed through the combination of PARPi or p21 depletion does not cause of PARPi sensitivity.

### Gap suppression in de novo models of PARPi resistance

If gaps underlie therapy response, then gap suppression should trigger PARPi resistance (**Figure 3A**). To determine whether gap suppression occurs in models of PARPi resistance, we analyzed cell lines that were initially sensitive to PARPi and then had gained PARPi resistance while propagated in tissue culture(Yazinski et al., 2017). We first analyzed BRCA1-deficient mouse ovarian tumor cell line, BR5 and its derived PARPi-resistant cell line, BR5-R1(Yazinski et al., 2017). We confirmed PARPi sensitivity of BR5 as compared to BR5-R1 and BRCA1 proficient T2 cells (Yazinski et al., 2017) (**Figure 3B)**. We observed that PARPi treatment induced greater replication tract lengthening and gap induction in BRCA1 deficient BR5 cells as compared to BRCA1 proficient T2 cells (**Figure 3C**). In contrast, PARPi resistant BR5-R1 cells did not gain longer tracts and PARPi induced gaps were modest unless co-treated with the ATR inhibitor (ATRi), VE-821, that re-establishes PARPi sensitivity (Yazinski et al., 2017) (**Figure 3B-D, S2A**). *De novo* PARPi resistance is also associated with BRCA reversion mutations that restore function (**Figure S2B)**. In the BRCA2 reversion cell clone C4-2 that derived from the parental BRCA2 mutant PEO1 ovarian cancer cell line (Sakai et al., 2009), we observed restored fork restraint and gap suppression (**Figure S2C,D)** indicating that gap suppression (GS), HR and/or FP could confer PARPi resistance.

**Figure 3:**
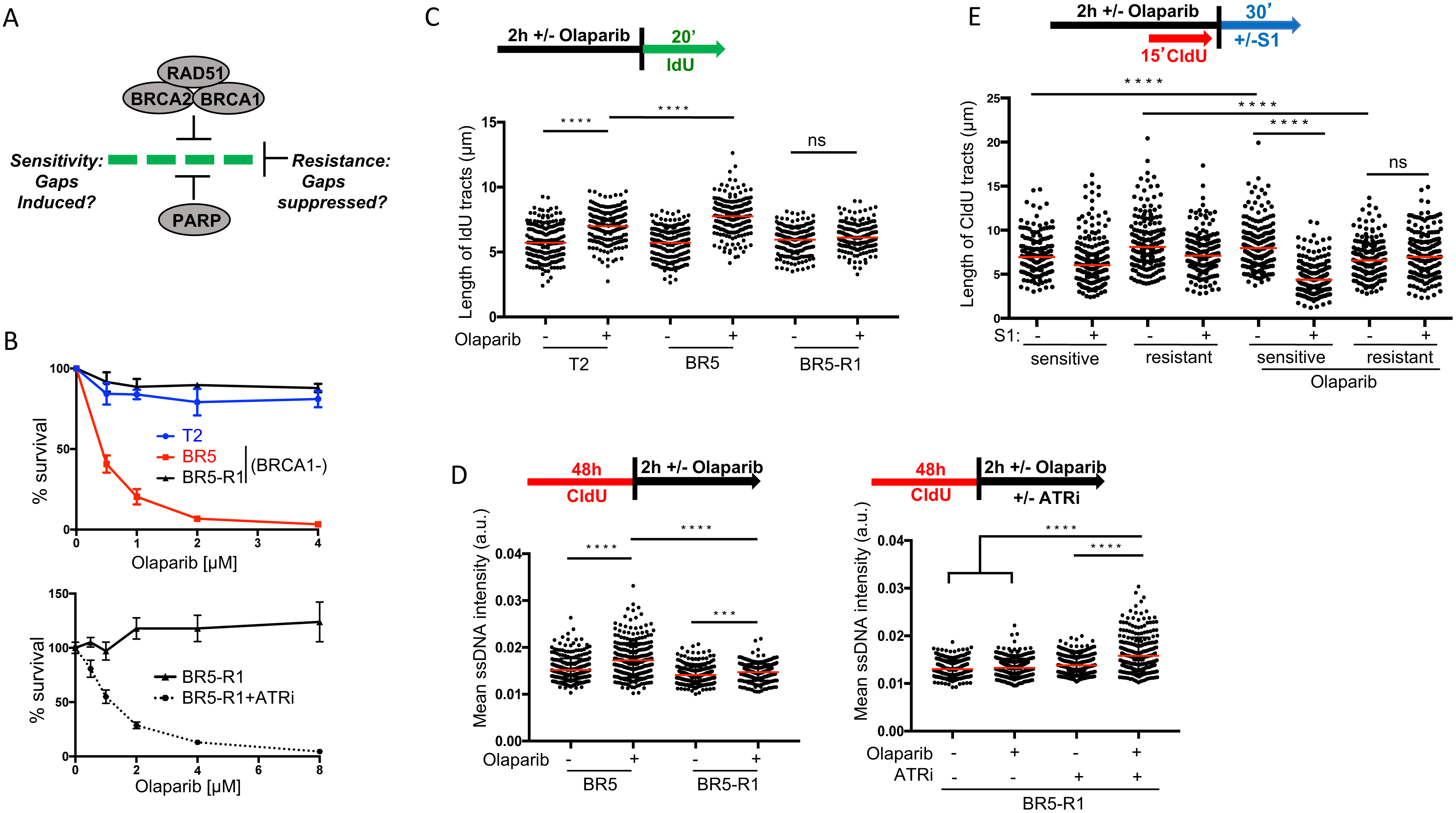
Gap suppression correlates with PARPi resistance. (A) Model depicting the hypothesis that gaps, prevented by BRCA-RAD51 and PARP, are the cause of PARPi sensitivity and gap suppression the mechanism of resistance. (B) Cell survival assays for T2, BR5 and BR5-R1 (BR5-derived resistant cells) cells under increasing concentrations of olaparib without or with ATR inhibitor (VE-821, 1μM). Data represent the mean percent ± SD of survival for each dot. (C) Schematic and quantification of the length of IdU tracts in indicated cells following olaparib treatment (10μM, 2 hours). Each dot represents one fiber; at least 200 fibers are quantified from two independent experiments. Red bars represent the median. Statistical analysis according to two tailed Mann-Whitney test. (D) Schematic and quantification of mean ssDNA intensity for BR5 and BR5-R1 cells following CldU pre-labeling and olaparib release (10μM, 2 hours), with or without ATR inhibitor (VE-821, 1μM). At least 200 cells are quantified from two independent experiments. Bars represent the mean ± SD. Statistical analysis according to t test. (E) Schematic and quantification of the length of IdU tracts with or without S1 nuclease incubation in indicated PDX samples and olaparib treatment (10μm, 2 hours). Each dot represents one fiber; at least 150 fibers are quantified. Red bars represent the median. Statistical analysis according to two tailed Mann-Whitney test. All p values are described in Statistical methods.

Next, we analyzed patient derived xenograft (PDX) that while initially sensitive to PARPi, gained resistance while propagated upon serial passage in mice. Tumor samples that were PARPi sensitive showed significant PARPi-induced lengthening of replication tracts (**Figure 3E**). Consistent with this lengthening reflecting replication with gaps, elongated tracts dramatically shortened upon S1-nuclease treatment (**Figure 3E**). In contrast, PARPi resistant tumor samples did not show PARPi-induced lengthening or S1 nuclease shortening (**Figure 3E**), indicating that gaps were not present. Collectively, these models of acquired PARPi resistance demonstrate a block to PARPi-induced acceleration that causes gaps.

### Gaps correlate with PARPi response better than HR or FP status

To further consider the relationship between gaps and PARPi response, we analyzed several well-established genetic models of PARPi resistance associated with or without HR and/or FP. In particular, HR and PARPi resistance are re-established in BRCA1 deficient cells by loss of the DNA repair protein 53BP1, which limits HR by blocking DNA end resection(Bouwman et al., 2010; Bunting et al., 2010). We confirmed that 53BP1 deletion in BRCA1 K/O RPE1 cells or 53BP1 depletion in BRCA1 deficient BR5 cells enhances PARPi resistance(Noordermeer et al., 2018) (**Figure 4A,B and S3A,B**). As compared to BRCA1 deficiency alone, loss of both BRCA1 and 53BP1 reduced PARPi-induced gaps (**Figure 4C, S3C**), suggesting that 53BP1 loss restores not only HR, but also suppresses gaps.

**Figure 4:**
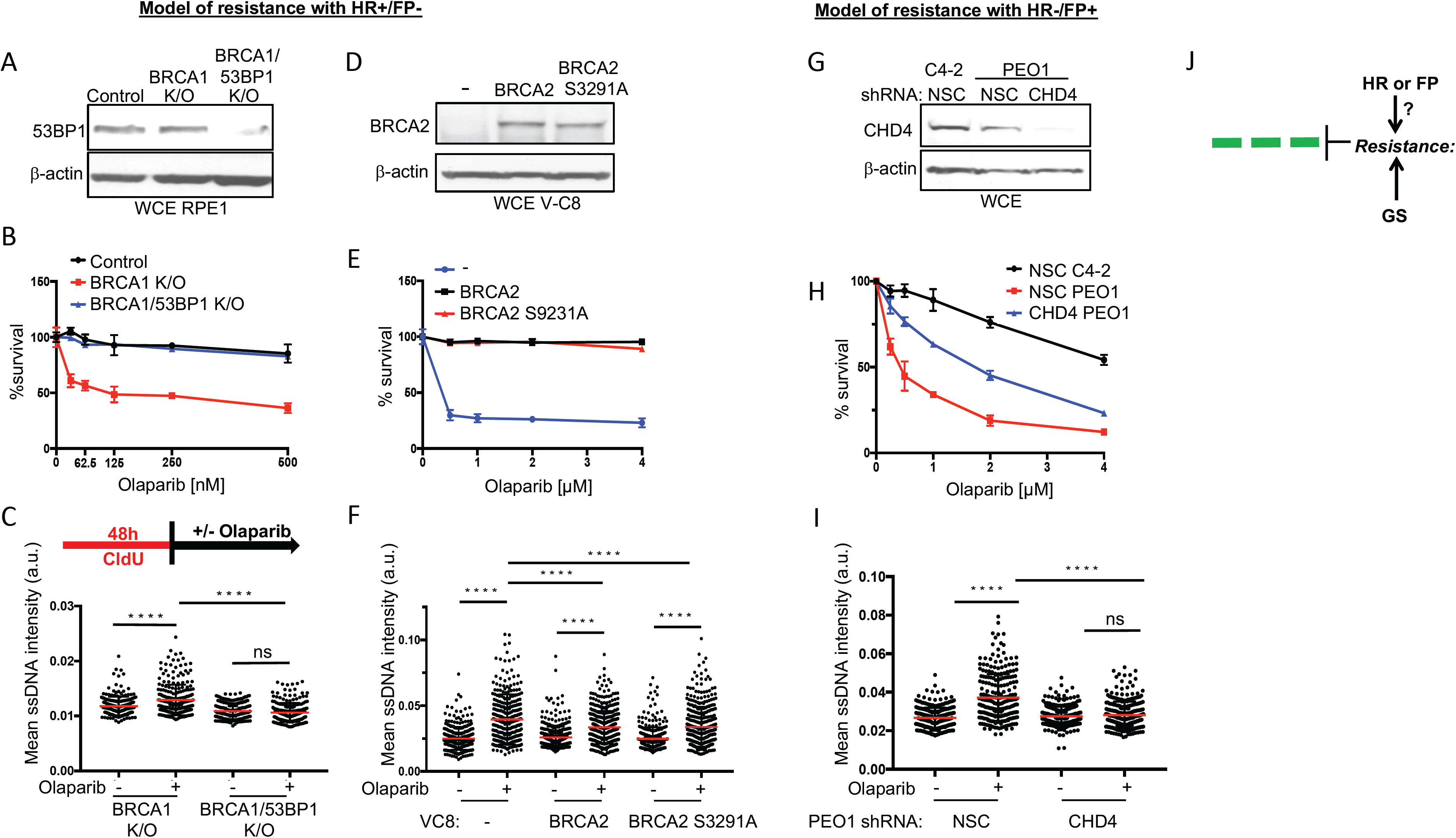
Gap Suppression and HR or FP correlate with PARPi resistance. (A) Western blot analysis with the indicated antibodies of lysates from Control, BRCA1 K/O and BRCA1/53BP1 K/O RPE1 cells. (B) Cell survival assays for Control, BRCA1 K/O and BRCA1/53BP1 K/O RPE1 cells under increasing concentrations of olaparib. Data represent the mean percent ± SD of survival for each dot. (C) Schematic and quantification of mean ssDNA intensity for indicated cell lines following CldU pre-labeling and olaparib release (10μM, 2 h). (D) Western blot analysis with the indicated antibodies of lysates from V-C8, V-C8+BRCA2 and V-C8+BRCA2 S3291A cells. (E) Cell survival assays for V-C8, V-C8+BRCA2 and V-C8+BRCA2 S3291A cells under increasing concentrations of olaparib. Data represent the mean percent ± SD of survival for each dot. (F) Quantification of mean ssDNA intensity for V-C8 and indicated cells following CldU pre-labeling and olaparib release (1μM, 4 hours). (G) Western blot analysis with the indicated antibodies of lysates from C4-2 and PEO1 cells expressing shRNA against NSC and CHD4. (H) Cell survival assays for C4-2 NSC, PEO1 NSC and PEO1 shCHD4 cells under increasing concentrations of olaparib. Data represent the mean percent ± SD of survival for each dot. (I) Quantification of mean ssDNA intensity for C4-2 and PEO1 cells expressing shRNA against NSC and CHD4 following CldU pre-labeling and olaparib release (1μM, 4 h). For (C), (F) and (I), at least 200 cells are quantified from two independent experiments. Bars represent the mean ± SD. Statistical analysis according to t test. All p values are described in Statistical methods. (J) Model indicating that GS co-occurs with either HR or FP confounding the PARPi resistance mechanism.

PARPi resistance is also re-established in BRCA2 mutant cells with or without FP. In particular, complementation of the BRCA2 mutant Chinese hamster cells, V-C8 with the C-terminal BRCA2 S3291A mutant that is defective in FP restores PARPi resistance similar to BRCA2 wild-type (**Figure 4D,E**)(Schlacher et al., 2011). Moreover, we observed that both PARPi resistant cell lines had significantly fewer gaps than the vector complemented PARPi sensitive V-C8 cells (**Figure 4F**), suggesting that resistance stemmed from gaps being suppressed below a toxicity threshold. In BRCA2 deficient cells, therapy resistance and FP are also enhanced by depletion of the chromatin remodeler, CHD4 (Guillemette et al., 2015; Ray Chaudhuri et al., 2016) Notably, CHD4 depletion not only enhanced PARPi resistance, but also dramatically suppressed PARPi-induced gaps as compared to control (**Figure 4G-I**). Based on these described models, it remains unclear if GS, HR or FP underlie the mechanism of PARPi resistance (**Figure 4J**). Moreover, in FANCJ null cells without gaps, and defective HR and FP (Peng et al., 2018),(Cantor et al., 2004; Levitus et al., 2005; Levran et al., 2005; Litman et al., 2005; Nath et al., 2017; Peng et al., 2018; Peng et al., 2006; Sawyer et al., 2015; Suhasini et al., 2013), it is possible that PARPi resistance reflects residual HR or FP activity.

To better understand the link between gaps and therapy response without the HR and FP confounding the interpretation of resistance mechanism, we flipped the model system to one in which despite HR and/or FP proficiency, cells remain PARPi sensitive allowing one to address if gaps remain (**Figure 5A**). HR is intact in the Fanconi anemia (FA) patient fibroblast cell line that has one wild-type and one mutant (T131P) RAD51 allele, but they remain therapy sensitive(Wang et al., 2015) as we confirmed as compared to a CRISPR corrected line (**Figure 5B**). While PARPi-induced gaps were observed in both wild-type and mutant cells, the mutant cell line had a higher level of gap induction and ratio of gaps (**Figure 5C)**. Conceivably, the FP defect in the FA cells could generate DSBs that overwhelm HR and sensitize to PARPi. FP can be restored in the RAD51 mutant FA cells by depletion of the RAD51 negative regulator, RADX(Bhat et al., 2018). Notably, despite restored FP upon RADX depletion, the RAD51 mutant FA cells remained PARPi sensitive (**Figure 5D,E and S3D**). Because PARPi gaps are observed and the gap ratio was elevated (**Figure 5F**), these findings suggest that HR and/or FP are insufficient for PARPi resistance when gaps exceed a toxicity threshold.

**Figure 5:**
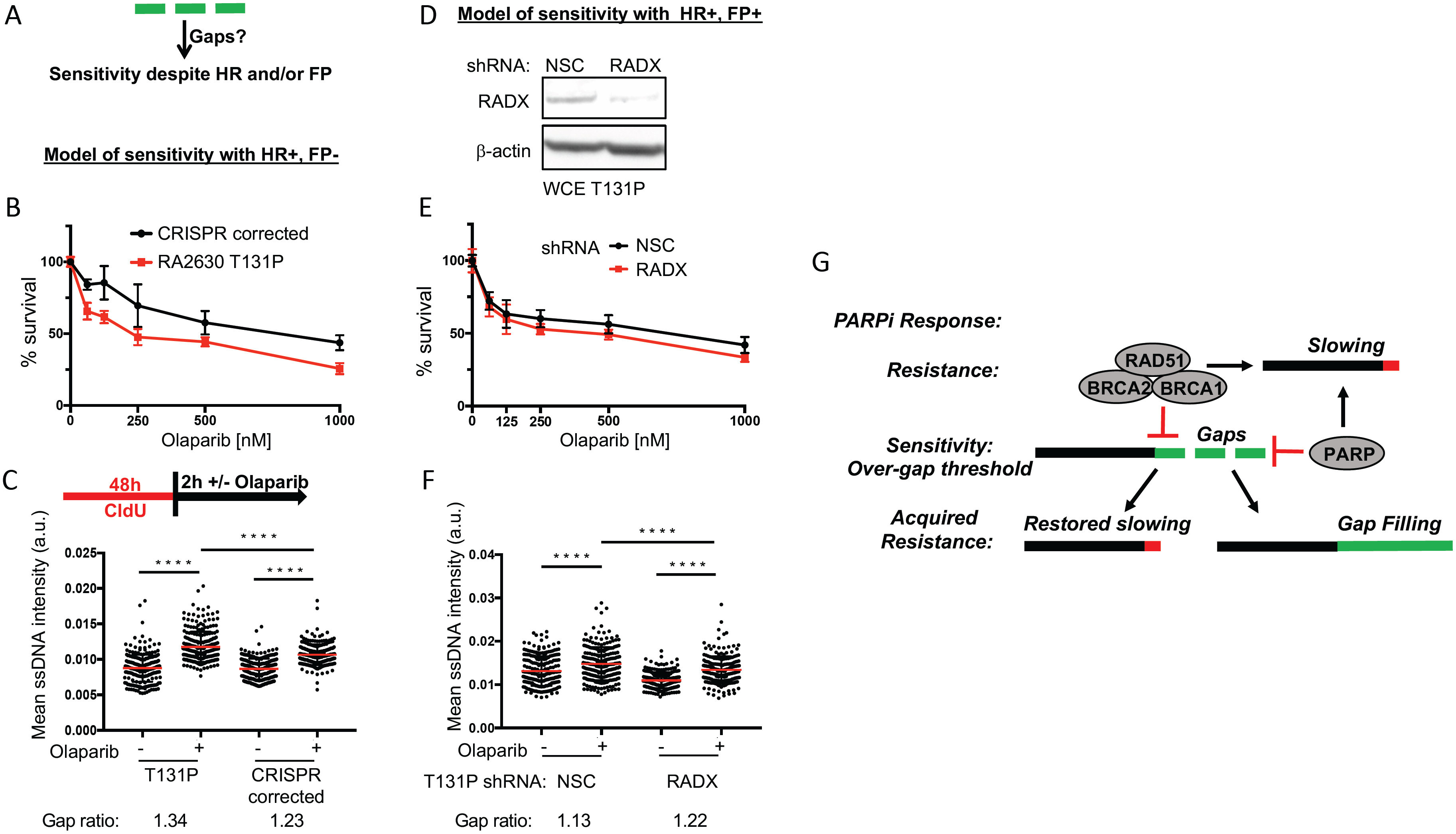
Gaps confer PARPi sensitivity despite HR and/or FP proficiency. (A) Could gaps sensitize cells despite HR or FP proficiency? (B) Cell survival assays for patient fibroblasts (RA2630) RAD51 T131P and RAD51 CRISPR corrected cells under increasing concentrations of olaparib. Data represent the mean percent ± SD of survival for each dot. (C) Schematic and quantification of mean ssDNA intensity for indicated cell lines following CldU pre-labeling and olaparib release (10μM, 2 h). The gap ratio is the fold change of ssDNA intensity from treated to untreated. (D) Western blot analysis with the indicated antibodies of lysates from patient fibroblasts (RA2630) RAD51 T131P cells expressing shRNA against non-silencing control (NSC) and RADX. (E) Cell survival assays for patient fibroblasts (RA2630) RAD51 T131P expressing shRNA against non-silencing control (NSC) and RADX under increasing concentrations of olaparib. Data represent the mean percent ± SD of survival for each dot. (F) Quantification of mean ssDNA intensity for indicated cell lines following CldU pre-labeling and olaparib release (10μM, 2 hours). For (C) and (F), at least 200 cells are quantified from two independent experiments. Bars represent the mean ± SD. Statistical analysis according to t test. All p values are described in Statistical methods. The gap ratio is the fold change of ssDNA intensity from treated to untreated. (G) Model summarizing that gaps are the determent of PARPi therapy response.

## DISSCUSSION

PARP1 function and loss-of-function has been linked to ssDNA(Leppard et al., 2003; Lonskaya et al., 2005; Maya-Mendoza et al., 2018; Ray Chaudhuri and Nussenzweig, 2017). However, ssDNA has been overlooked as the sensitizing lesion in BRCA deficient cells in favor of DNA double stranded breaks. Challenging this break-induced model of therapy sensitivity, PARPi do not initially generate DNA breaks or pause DNA replication forks. Instead, PARPi dysregulates DNA replication as fork reversal and restraint are disrupted (Berti et al., 2013; Maya-Mendoza et al., 2018; Ray Chaudhuri et al., 2012; Sugimura et al., 2008; Zellweger et al., 2015). Therefore, the mechanism of action of PARPi-induced killing of BRCA deficient cells was in question. As clinical interventions for PARPi resistance are lacking, understanding the sensitizing lesion is of critical importance.

Here, we provide evidence that PARPi leads to dysregulated replication with the induction of ssDNA gaps that accumulate more excessively in BRCA deficient immortalized cells, human and mouse cancer cell lines, and patient tumors. Moreover, PARPi-induced gaps are restricted in FANCJ deficient cells that are not sensitive to PARPi. In BRCA1 deficient cells, PARPi-induced gaps are suppressed in both *de novo* and genetic models of PARPi resistance. Further indicating gaps as the sensitizing lesion, models of restored or defective HR or FP reveal that despite differences in proficiency, sensitivity correlated with gap induction and resistance with gap suppression (**Figure 5G**). Our data also highlight that dysregulated or accelerated replication is not the cause of PARPi sensitivity. Together, these findings reveal that gaps as opposed to loss of HR or FP proficiency more directly predicts PARPi response (**Table 1)**.

**Table1.**
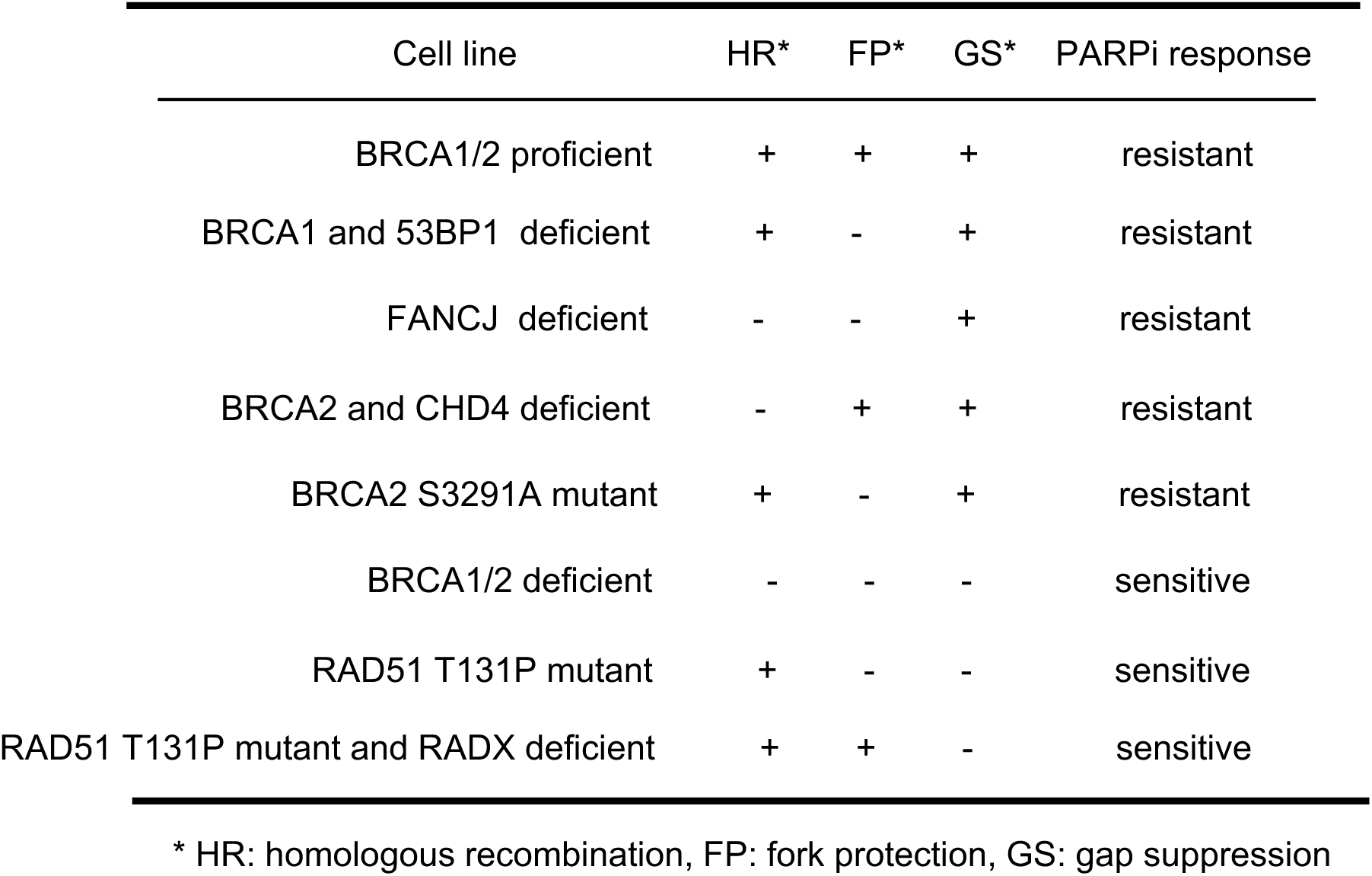
Hallmarks of PARPi response.

Rather our findings indicate that toxicity results from gaps that exceed a safe threshold. We envision that the toxicity threshold is more readily exceeded in BRCA deficient cells due to intrinsic defects in coordinating RAD51 replication restraint and gap avoidance functions ((Hashimoto et al., 2010; Kolinjivadi et al., 2017b; Zellweger et al., 2015)). Indeed, we demonstrate that in response to a range of drugs including cisplatin, BRCA deficiency interferes with replication restraint and gaps develop (Panzarino et al., 2019). Thus, we propose that HR deficiency is not the root cause of therapy response or for that matter “BRCAness”, but rather gaps are the key factor.

In cells defective in restraining replication in response to stress, PARPi treatment could lead to unrestrained replication that proceeds until forks confront PARP-trapped proteins or other intrinsic obstacles that physically block the pre-existing replication fork. Gaps could form as replication re-initiates ahead of the block by re-priming replication. The extensive regions of under-replication could drive RPA exhaustion and replication catastrophe(Toledo et al., 2013). Alternatively, given that ssDNA marks apoptotic cells and ssDNA induces apoptosis(Chen et al., 1997; Frankfurt et al., 1996; Gagna et al., 2000; Michiue et al., 2008; Nur et al., 2003), the accumulation of an excessive amount of ssDNA could be the single driver of cell killing. Our findings further indicate that PARPi resistance involves gap suppression by either a block to fork elongation or by the acquisition of gap filling activity such as mediated by translesion synthesis (TLS). In support of this latter point, PARPi resistance in BRCA2 deficient cells is achieved by loss of CHD4 that elevates TLS(Guillemette et al., 2015). Moreover, p21 is a TLS inhibitor(Avkin et al., 2006) and its loss lengthens tracts without gaps. Notably, 53BP1 loss uniquely rescues BRCA1 deficient cells and CHD4 loss uniquely rescues BRCA2 deficient cells(Bouwman et al., 2010; Bunting et al., 2010; Guillemette et al., 2015; Ray Chaudhuri et al., 2016). Thus, the mechanism of gap suppression is distinct in BRCA backgrounds for reasons that remain to be fully understood.

While we cannot exclude the possibility that eventual PARPi-induced fork degradation and/or breaks contribute to sensitizing BRCA deficient cells, we present a series of separation-of-function models that indicate a greater correlation between gap induction and therapy response than loss of either HR or FP. Moreover, if DSBs were the sensitizing lesion, logically HR proficient cells should repair PARPi-induced DSBs and show PARPi resistance. However, the HR proficient FA cell line is sensitive to PARPi (Wang et al., 2015) even when FP is also restored. These findings indicate that HR and FP are not sufficient to confer PARPi resistance if gaps exceed a toxicity threshold. Indeed, the rationale for limiting PARPi to HR defective cancers is already in question as recent clinical trials found significant clinical benefit across ovarian cancer patients regardless of BRCA status (Ledermann and Pujade-Lauraine, 2019). Moreover, our study indicates that to further improve PARPi progression free, overall survival or in maintenance therapy (Miller et al., 2019), it will be critical to target gap suppression pathways. Collectively, these findings highlight the importance of considering ssDNA gaps as a critical biomarker and determinant of therapy response.

## FIGURE LEGENDS

**Figure S1:**
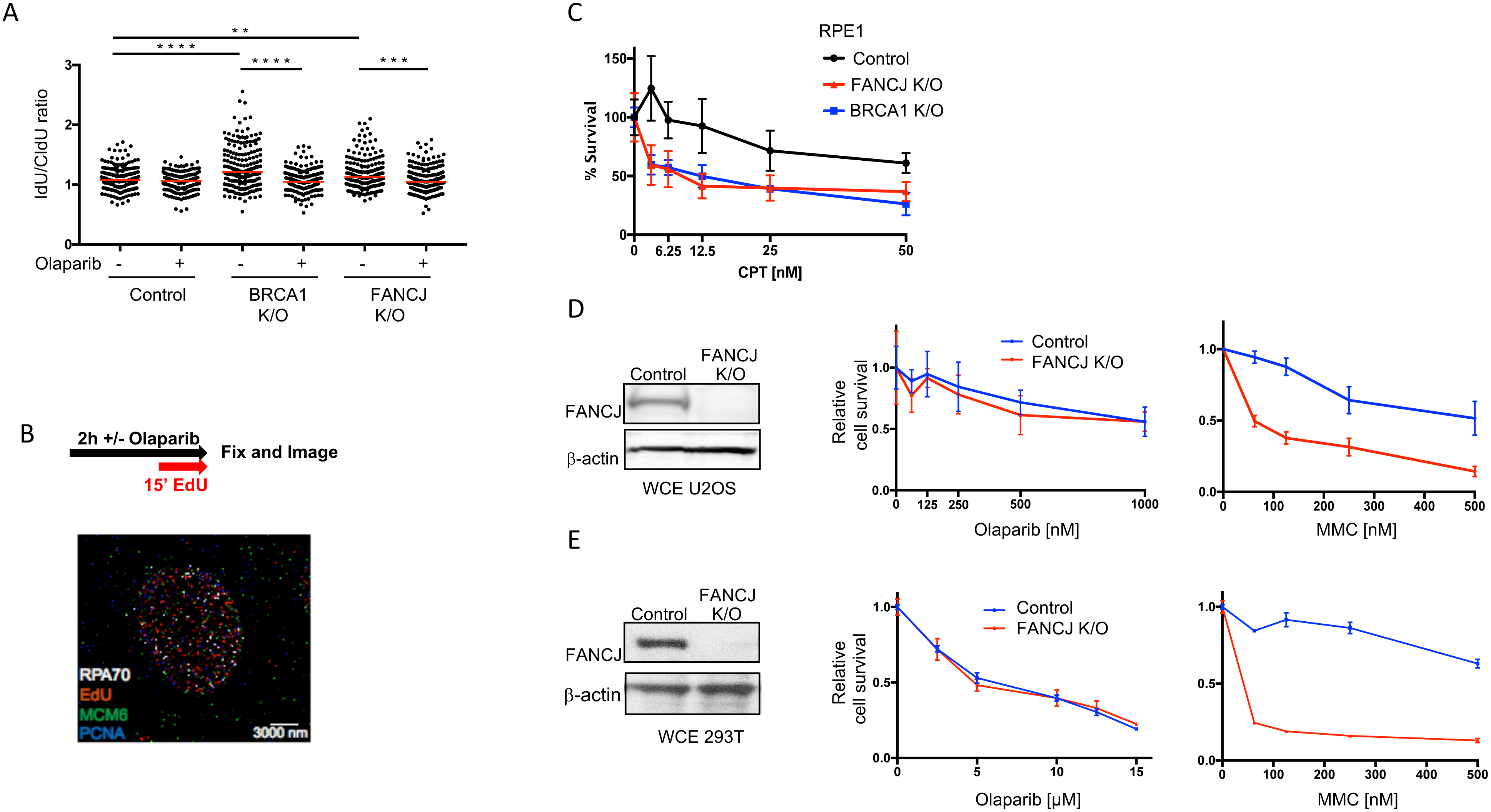
Status of fork asymmetry and responses to stress. (A) Quantification of the ratio of IdU/CldU in indicated cells following olaparib treatment (10μM, 2 hours). Each dot represents one fiber; at least 200 fibers are quantified from two independent experiments. Red bars represent the median. Statistical analysis according to two tailed Mann-Whitney test. (B) Experimental scheme for STORM imaging of replication forks and representative super-resolution image of a single U2OS nucleus labeled for nascent DNA (using EdU, red), RPA70 (grey), MCM6 (green) and PCNA (blue). Scale bar, 3000nm. (C) Cell survival assays for Control, FANCJ K/O and BRCA1 K/O RPE1 cells under increasing concentrations of camptothecin (CPT). Data represent the mean percent ± SD of survival for each dot. (D) and (E) Western blot analysis with the indicated antibodies of lysates from Control, FANCJ K/O U2OS cells (C) and 293T cells (D). Cell survival assays for indicated cells under increasing concentrations of olaparib and mitomycin C (MMC). Data represent the mean percent ± SD of survival for each dot.

**Figure S2:**
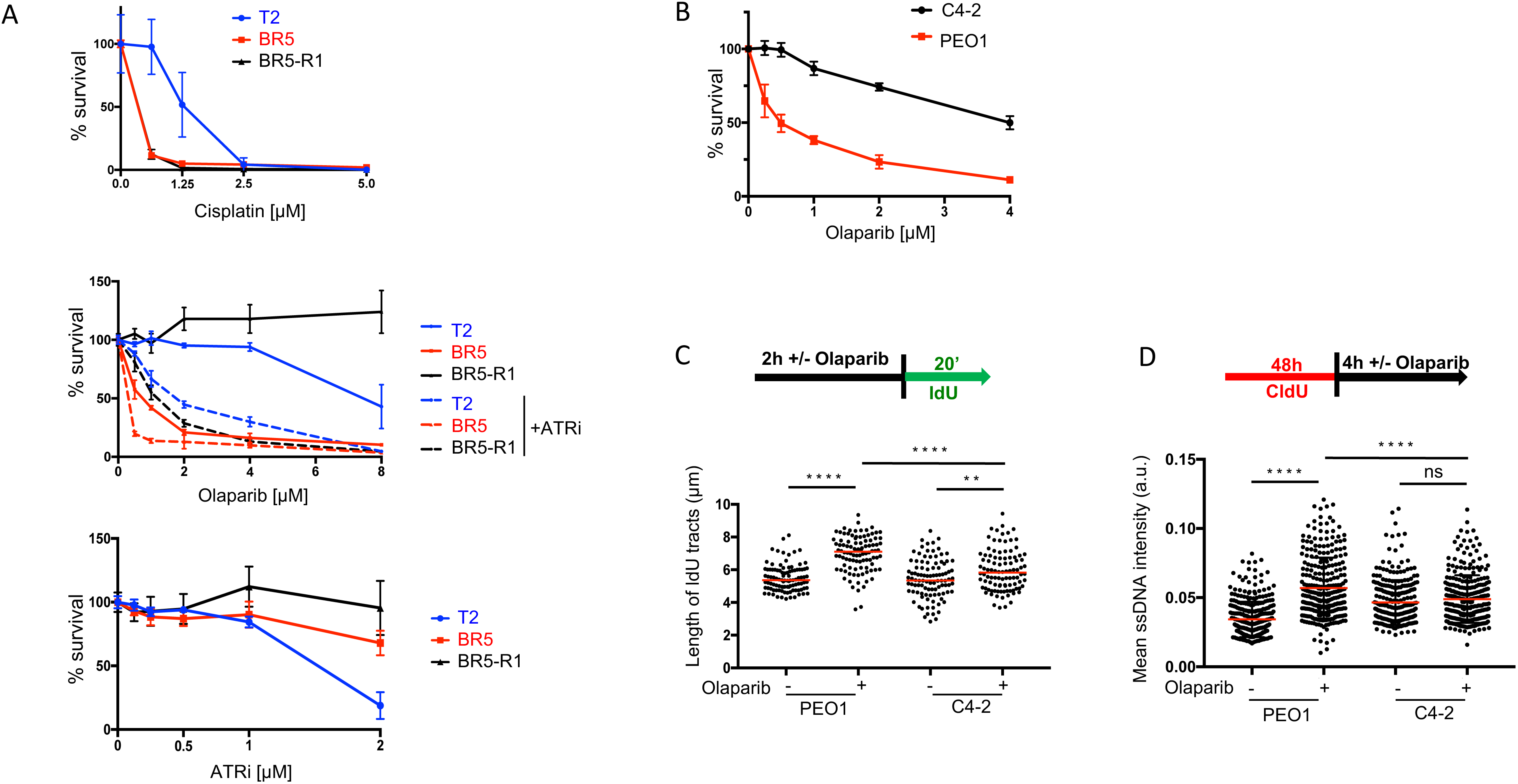
Sensitivity, fork acceleration and gaps in response to stress. (A) Cell survival assays for T2, BR5 and BR5-R1 cells under increasing concentrations of olaparib with ATR inhibitor (VE-821, 1μM) and ATR inhibitor alone. Data represent the mean percent ± SD of survival for each dot. (B) Cell survival assays for C4-2 and PEO1 cells under increasing concentrations of Olaparib. Data represent the mean percent ± SD of survival for each dot. (C) Schematic and quantification of the length of IdU tracts in PEO1 and C4-2 cells following olaparib treatment (10μM, 2 hours). Each dot represents one fiber, at least 100 fibers are quantified. Red bars represent the median. Statistical analysis according to two tailed Mann-Whitney test. (D) Schematic and quantification of mean ssDNA intensity for indicated cell lines following CldU pre-labeling and olaparib release (1μM, 4 hours). At least 200 cells are quantified from two independent experiments. Bars represent the mean ± SD. Statistical analysis according to t test. All p values are described in Statistical methods.

**Figure S3:**
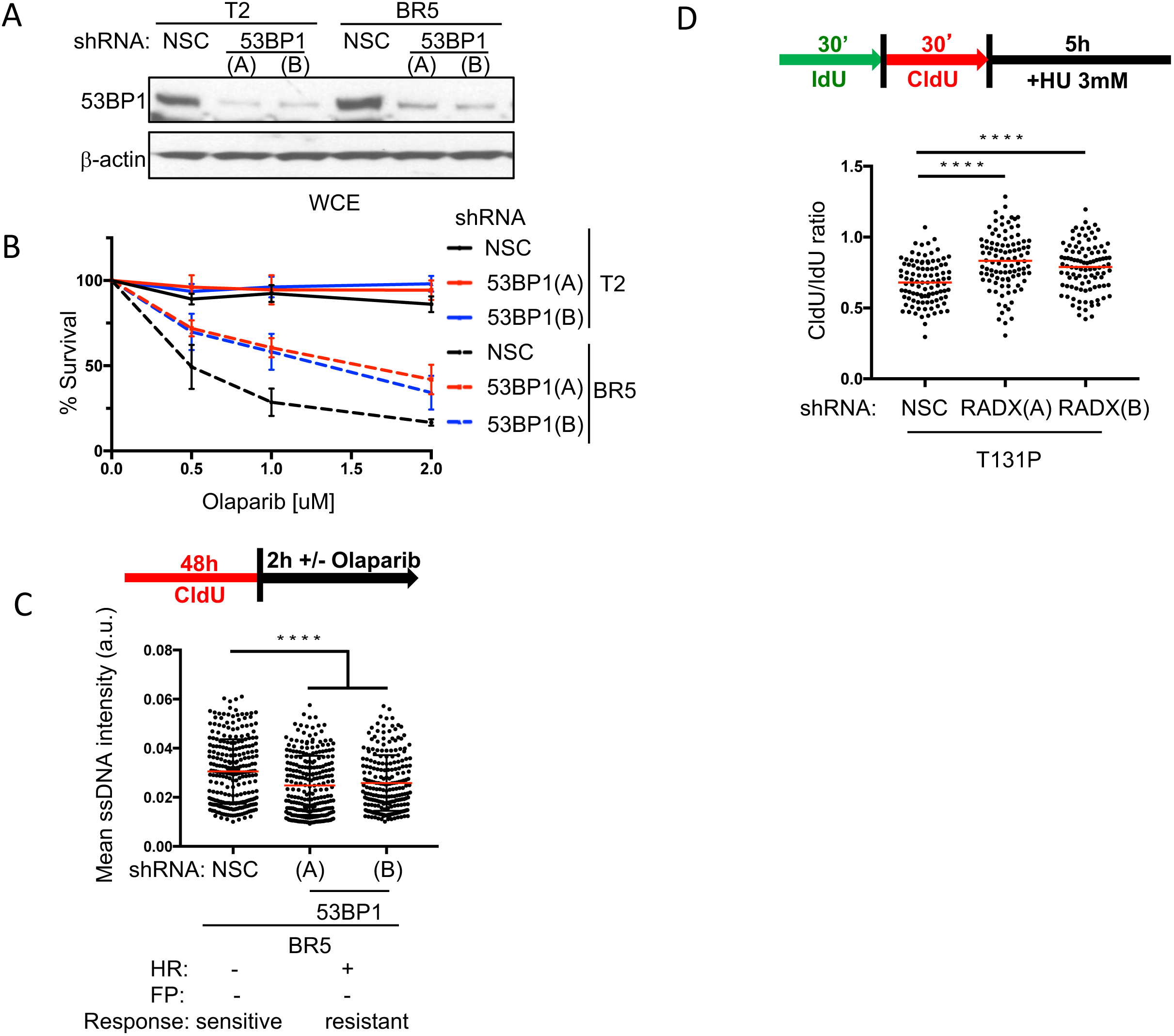
Analysis of rescue of BRCA-RAD51 deficiency. (A) Western blot analysis with the indicated antibodies of lysates from T2 and BR5 cells expressing shRNA against NSC, 53BP1(A) and 53BP1(B). (B) Cell survival assays for indicated cells under increasing concentrations of olaparib. Data represent the mean percent ± SD of survival for each dot. (C) Quantification of mean ssDNA intensity for BR5 cells expressing shRNA against NSC, 53BP1(A) and 53BP1(B) following CldU pre-labeling and olaparib release (10μM, 2 hours). At least 200 cells are quantified from two independent experiments. Bars represent the mean ± SD. Statistical analysis according to t test. (D) Schematic and quantification of the CldU/IdU ratio after 3mM HU treatment for 5 h in patient fibroblasts (RA2630) RAD51 T131P expressing shRNA against non-silencing control (NSC) and RADX. Each dot represents one fiber. For each analysis, at least 100 fibers are quantified. Red bars represent the median. Statistical analysis according to two-tailed Mann-Whitney test. All p values are described in Statistical methods.

## METHODS

### Cell lines and gene editing

Human RPE1-hTERT cell lines were grown in DMEM+GlutaMAX-I (Gibco, 10569) supplemented with 10% FBS and 1% Pen Strep (100 U/ml). U2OS, 293T, PEO1 and C4-2 cell lines were grown in DMEM (Gibco, 11965) supplemented with 10% fetal bovine serum (FBS) and 1% Pen Strep (100 U/ml). T2, BR5, BR5-R1 cell lines were cultured in DMEM (CORNING cellgro, 15-017-CV) with 10% FBS, penicillin and streptomycin (100 U/ml each), and 1% L-glutamine(Yazinski et al., 2017). The resistant cell line BR5-R1 was maintained in 1μM olaparib. FA Patient fibroblasts (RA2630 T131P and CRISPR corrected clone 3-39) were cultured in DMEM (Gibco, 11965) supplemented with 10% FBS, 1% Pen Strep (100 U/ml), 1% GlutaMAX-I, 1% MEM NEAA and 1% Sodium pyruvate(Wang et al., 2015). All the cell lines were cultured at 37 °C, 5% CO_2_. The generation of RPE1-hTERT TP53−/− BRCA1 K/O and BRCA1/53BP1 double K/O Cas9 cells were described elsewhere(Noordermeer et al., 2018). FANCJ gene knockout in RPE1-hTERT TP53−/− Cas9 cells was introduced by using two synthesized gRNAs (gRNA #1: GGGTCGAGGAAAGGTAACGG, gRNA #2: GGCAATCACCACACCCTTCA) and the protocol from IDT technology. For generating FANCJ gene knockout in RPE1-hTERT TP53−/− Cas9 cells, 10 μl tracrRNA (100 μM) and 10 μl 20 nt crRNA (100 μM) were annealed in 80 μl nuclease free duplex buffer (IDT#11-05-01-03) to form a 10 μM gRNA solution. The crRNAs were designed using CRISPR design tools of Benchling. In brief, 100.000 cells were seeded out the day before transfection in 12-well dish in 1 mL medium. 30 minutes prior to transfection the medium was replaced with 750 μl of medium. The 3 μl of 10 μM for each gRNA was added to optiMEM (Life Technologies) to final volume of 125 μl and incubated briefly at RT. Then 6 μl Lipofectamine RNAiMAX (Invitrogen) was added and supplemented to a total volume of 250 μl with optiMEM. Mixture was incubated at RT for 20 min before adding to the cells. The next day the medium was changed, and 2 days after transfection the cells were serial diluted in 96 well plates to obtain single clones. 2 weeks later, the cell clones were passages for new 96 wells, and screened for full loss of both BRIP1 alleles using PCR approach (PCR primer set flanking 5’ end (PCR#78: TTCCATTGGATGCCGAAGT, PCR#79: CGCTCAAAGGAGGTAAGGATAG), and one primer set flanking the full-length gene (PCR#78 + PCR#43, CCACAACACGTCGGGATTAT). Successful KO clones were validated by WB. U2OS FANCJ K/O and 293T FANCJ K/O cells were generated and maintained as previously described(Peng et al., 2018).

### Immunoblotting and antibodies

Cells were harvested, lysed and processed for Western blot analysis as described previously using 150mM NETN lysis buffer (20mM Tris (pH 8.0), 150mM NaCl, 1mM EDTA, 0.5% NP-40, 1mM phenylmethylsulfonyl fluoride, 10mg/ml leupeptin, 10mg/ml aprotinin). Proteins were separated using SDS–PAGE and electro-transferred to nitrocellulose membranes. Membranes were blocked in 5% not fat dry milk (NFDM) phosphate-buffered saline (PBS)/Tween 20 and incubated with primary antibodies for overnight at 4°C. Antibodies for Western blot analysis included anti-β-actin (Sigma), anti-FANCJ (E67), anti-BRCA1 (Cell Signaling Technology), anti-p21 (Cell Signaling Technology), anti-53BP1(Novus Biological) and anti-CHD4 (Abcam). Membranes were washed, incubated with corresponding horseradish peroxidase-linked secondary antibodies (Amersham, GE Healthcare) for 1h at room temperature and detected by chemiluminescence imaging system (Bio-Rad).

### RNA interference

U2OS cells were reverse transfected using RNAiMAX transfection reagent (Life Technologies 13778150) and siRNA targeting FANCJ/BRIP1 (QIAGEN SI03110723), BRCA1 (QIAGEN SI00299495), or scrambled negative control (ORIGENE SR30004) in 6-well plates for 48 h before super-resolution microscopy analysis. Stably transduced cells were generated by infection with pLK 0.1 vectors containing shRNAs against non-silencing control (NSC) or one of the shRNAs against corresponding genes: p21(CDKN1A) includes (A) 5′-TAAGGCAGAAGATGTAGAGCG-3′, (B) 5′-AAAGTCGAAGTTCCATCGCTC-3′; 53BP1(TP53BP1) includes (A) 5′-AAACCAGTAAGACCAAGTATC-3′, (B) 5′-AATCAATACTAATCACACTGG-3′; CHD4 includes 5’-AATTCATAGGATGTCAGCAGC-3’; RADX(CXorf57) includes 5’ ATTTCCGTGGAATACTTTCAG-3’. The sequence information was obtained from Dharmacon website (https://dharmacon.horizondiscovery.com), and the shRNAs were obtained from the University of Massachusetts Medical School (UMMS) shRNA core facility. Cells were selected by puromycin for 3-5 days before experiments were carried out.

### Drugs and reagents

The following drugs were used in the course of this study: PARP inhibitor olaparib (AZD-2281, SelleckChem), cisplatin (Sigma-Aldrich), camptothecin (Sigma-Aldrich), ATR inhibitor (VE-821, SelleckChem). Reagents used including 5-chloro-2′-deoxyuridine (CldU) and 5-Iodo-2′-deoxyuridine (IdU) were obtained from Sigma-Aldrich. Concentration and duration of treatment are indicated in the corresponding figures and sections.

### Immunofluorescence for ssDNA

Cells were grown on coverslips in 10 μM CldU for 48 h before the indicated treatment in figures without CldU. After treatment, cells were washed with PBS and pre-extracted by 0.5% Triton X-100 made in phosphate-buffered saline (PBS) on ice. Cells were then fixed using 4% Formalin for 15 min at RT, and then permeabilized by 0.5% Triton X-100 in PBS again. Permeabilized cells were then incubated with primary antibodies against CldU (Abcam 6326) at 37°C for 1h. Cells were washed and incubated with secondary antibodies (Alexa Fluor 594) for 1 h at room temperature. After washing, cover slips were mounted onto glass slides using Vectashield mounting medium containing DAPI (Vector Laboratories). Images were collected by fluorescence microscopy (Axioplan 2 and Axio Observer, Zeiss) at a constant exposure time in each experiment. Representative images were processed by ImageJ software. Mean intensities of ssDNA in each nucleus were measured with Cell Profiler software version 3.1.5 from Broad Institute.

### DNA fiber assay and S1 nuclease analysis

These assays were performed as previously described(Peng et al., 2018). Cells were labeled by sequential incorporation of two different nucleotide analogs, IdU and CldU, into nascent DNA strands for the indicated time and conditions. After nucleotide analogs were incorporated in vivo, the cells were collected, washed, spotted, and lysed on positively charged microscope slides by 7.5 mL spreading buffer for 8 min at room temperature. For experiments with the ssDNA-specific endonuclease S1, after the CldU pulse, cells were treated with CSK100 buffer for 10 min at room temperature, then incubated with S1 nuclease buffer with or without 20 U/mL S1 nuclease (Invitrogen, 18001-016) for 30 min at 37 C. The cells were then scraped in PBS + 0.1% BSA and centrifuged at 7,000 rpm for 5 min at 4 C. Cell pellets were resuspended at 1,500 cells/mL and lysed with lysis solution on slides. Individual DNA fibers were released and spread by tilting the slides at 45 degrees. After air-drying, fibers were fixed by 3:1 methanol/acetic acid at room temperature for 3 min. After air-drying again, fibers were rehydrated in PBS, denatured with 2.5 M HCl for 30 min, washed with PBS, and blocked with blocking buffer for 1 hr. Next, slides were incubated for 2.5 hr with primary antibodies for (IdU, Becton Dickinson 347580; CldU, Abcam 6326) diluted in blocking buffer, washed several times in PBS, and then incubated with secondary antibodies (IdU, goat anti-mouse, Alexa 488; CldU, goat anti-rat, Alexa Fluor 594) in blocking buffer for 1 hr. After washing and air-drying, slides were mounted with Prolong (Invitrogen, P36930). Finally, visualization of green and/or red signals by fluorescence microscopy (Axioplan 2 imaging, Zeiss) provided information about the active replication directionality at the single-molecule level.

### Viability assays

Cells were seeded onto 96-well plates (500 cells per well, performed in triplicate for each experiment group) and incubated overnight. Patient fibroblasts were seeded onto 6-well plates (5000 cells per well, performed in duplicate for each experiment group). The next day, cells were treated with increasing doses of drugs indicated in corresponding figures and maintained in complete media for 5 to 7 days. Percentage survival was measured either photometrically using a CellTiter-Glo 2.0 viability assay (Promega) in a microplate reader (Beckman Coulter DTX 880 Multimode Detector) or by manual cell counting.

### STROM analysis

For super resolution imaging experiments, cells were trypsinized and seeded on glass coverslips in six-well plates in low density. siRNA transfection, drug treatment, and EdU incorporation were performed directly on cells on coverslips. We used an optimized pre-extraction and fixation protocol for our immunofluorescence experiments in order to clearly visualize chromatin-bound nuclear fraction of cells and to minimize nonspecific antibody labeling from the cytoplasm and non-chromatin bound proteins that could significantly increase noise for image analysis. Cells were permeabilized with 0.5% Triton X-100 in ice-cold CSK buffer (10 mM Hepes, 300 mM Sucrose, 100 mM NaCl, 3 mM MgCl_2_, and 0.5% Triton X-100, pH = 7.4) in RT for 10 minutes and fixed with 4% paraformaldehyde (Electron Microscopy Sciences 15714) in RT for 30 minutes. Following fixation, cells were washed twice with PBS and blocked with blocking buffer (2% glycine, 2% BSA, 0.2% gelatin, and 50 mM NH_4_Cl in PBS). For nascent DNA detection, cells were pulse-labeled with 10 µM EdU (ThermoFisher A10044), a thymidine analogue, 15 minutes before permeabilization and fixation so that it would be incorporated into nascent DNA during replication in S-phase cells. After fixation, EdU was tagged with Alexa Fluor 647 picolyl azide through click reaction (Click-iT chemistry, ThermoFisher, C10640). The cells were blocked with blocking buffer at least overnight at 4°C. Before imaging, the samples were stained with primary antibodies against rb-RPA70 (Abcam ab79398), rb-MCM6 (conjugated to AF568, Abcam ab211916), and ms-PCNA (Santa cruz sc-56) in blocking buffer for 1h at RT, then secondary antibodies (goat anti-mouse AF488, Invitrogen A11029; goat anti-rabbit AF 750, Invitrogen A21039) in blocking buffer for 30minutes at RT. Super resolution imaging and other related processes were described before(Yin and Rothenberg, 2016).

### PDX studies

PNX0204 was derived at Fox Chase Cancer Center under IRB and IACUC approved protocols. PDX tumors were grown in NOD.Cg-Prkdcscid Il2rgtm1Wjl/SzJ (NSG) mice. Resistant PDX tumors were obtained from mice after tumors progressed on serial treatments of olaparib. The tumors were harvested at approximately 500 mm3 and dissociated in 0.2% collagenase, 0.33 mg/ml dispase solution for 3h at 37°C. The dissociated cells were maintained at 37°C in RPMI1640 + 10% FBS and used for DNA fiber assays within 24h of tumor extrackion. DNA fiber assay and S1 nuclease analysis were performed as described above.

### Statistical methods

Statistical differences in DNA fiber assays, immunofluorescence, and the alkaline BrdU comet assay were determined using unpaired t test or two-tailed Mann-Whitney test. Statistical analysis was performed using GraphPad Prism (Version 7.0). In all cases, ns: not significant (p > 0.01), **p < 0.01, ***p < 0.001, and ****p < 0.0001.

## ACKNOWLEDGEMENTS

We thank the members of the Cantor laboratory for helpful discussions. We thank Dr. Daniel Durocher for RPE1 cell lines including control, BRCA1 K/O and BRCA1/53BP1 K/O, Lee Zou for the T2, BR5, BR5-R1 cells, Agata Smogorzewska for the FA patient and CRISPR corrected cells, Toshi Taniguchi for the PEO1 and C4-2 cells and Maria Jasin for the VC-8 and derived cells. This work was supported by R01 CA176166-01A1 (Cantor), R01 CA214799 and OC130212 (Johnson) as well as charitable contributions from Mr. and Mrs. Edward T. Vitone, Jr. and the Lipp Family Foundation. We thank Drs. Igor Astutarov and Vladimir Khazak for establishing and sharing PNX0204.

## AUTHOR CONTRIBUTIONS

S.C. and K.C. designed the experiments. K.C., A.N.K., M.P., C.T.W.L., S.N., J.K., N.J.P., J.C. and M.B. performed the experiments. K.C. and M.P. analyzed the data. S.C. and K.C. wrote the manuscript. S.C., J.J., N.J. and, E.R. supervised the research.

